# CpgD is a phosphoglycerate cytidylyltransferase required for ceramide diphosphoglycerate synthesis

**DOI:** 10.1101/2025.01.09.632243

**Authors:** Tanisha Dhakephalkar, Ziqiang Guan, Eric A. Klein

## Abstract

Lipopolysaccharide (LPS) is essential in most Gram-negative bacteria, but mutants of several species have been isolated that can survive in its absence. *Caulobacter crescentus* viability in the absence of LPS is partially dependent on the anionic sphingolipid ceramide diphosphoglycerate (CPG2). Genetic analyses showed that *ccna_01210*, which encodes a nucleotidyltransferase, is required for CPG2 production. Using purified recombinant protein, we determined that CCNA_01210 (CpgD) is a phosphoglycerate cytidylyltransferase which uses CTP and 3-phosphoglycerate to produce CDP-glycerate, which we hypothesize is the phosphoglycerate donor for CPG2 synthesis. CpgD had optimum activity at pH 7.5-8 in the presence of magnesium. CpgD exhibited Michaelis-Menten kinetics with respect to 3-phosphoglycerate (Km,app = 10.9 ± 0.7 mM; Vmax,app = 0.72 ± 0.02 µmol/min/mg enzyme) and CTP (Km,app = 4.8 ± 0.9 mM; Vmax,app = 0.44 ± 0.03 µmol/min/mg enzyme). The characterization of this enzyme uncovers another piece of the pathway towards CPG2 synthesis.

## Introduction

The outer leaflet of the outer membrane of the cell envelope of Gram-negative bacteria is primarily composed of lipopolysaccharide (LPS) (1) which provides the first line of defense against environmental stress and hydrophobic antimicrobial drugs (2). LPS is essential in most Gram-negative species, but LPS-deficient mutants have been isolated from *Acinetobacter baumanii*, *Neisseria menengitidis*, *Moraxella catarrhalis*, and *Caulobacter crescentus* (3–6). While the mechanisms underlying survival in the absence of LPS appear to be species-specific, we previously showed that *C. crescentus* produces the anionic sphingolipid (SL) ceramide polyphosphoglycerate (CPG2) which is critical for viability upon LPS depletion (6).

SLs are built upon a ceramide backbone which consists of a sphingoid base and an N-linked fatty acid. Among the characterized bacterial SLs, these lipids can differ in acyl chain length, saturation, and branching, and contain a variety of headgroups. SLs play diverse roles in bacterial physiology including host-microbe interactions (7–9), defense against bacteriophages (10), and sporulation (11). The headgroup of CPG2, which supports survival in the absence of LPS, consists of a diphosphoglycerate moiety. Our previous genetic studies have identified at least four genes that are involved in the synthesis of CPG2 (6, 12), but their specific enzymatic activities remain largely unknown. Analysis of deletion mutants shows that *ccna_01218* (*cpgB*) and *ccna_01219* (*cpgC*) are required for adding the first phosphoglycerate to make ceramide phosphoglycerate (CPG), whereas *ccna_01217* (*cpgA*) and *ccna_01210* are involved in the addition of the second phosphoglycerate to produce CPG2.

In a previous report, we demonstrated that CpgB is a ceramide kinase which phosphorylates ceramide to produce ceramide 1-phosphate (C1P) (13). In the current study, we investigated the function of CCNA_01210 in producing CPG2. CCNA_01210 is annotated as a nucleotidyltransferase protein and it shares predicted structural homology with cytidylyltransferases which use CTP and sugar-phosphate substrates to produce CDP-sugars. The *Bacillus subtilis* cytidylyltransferase protein TagD is involved in wall teichoic acid (WTA) synthesis where it catalyzes the transfer of glycerol-3-phosphate to CTP, forming CDP-glycerol (14). TagF then uses CDP-glycerol as a substrate to transfer phosphoglycerol to its teichoic acid membrane acceptor (15). This modification of WTA has strong similarity to CPG2 leading to our hypothesis that CCNA_01210 uses CTP and 3-phosphoglycerate to form CDP-glycerate, which would later be used as a substrate to add a phosphoglycerate onto CPG to form CPG2. In this study, we used purified recombinant CCNA_01210 to characterize its enzymatic activity and confirmed its function as a CDP-glycerate producing cytidylyltransferase.

## Results

### CCNA_01210 is required for CPG2 production

*C. crescentus* synthesizes two novel anionic sphingolipids, CPG and CPG2 (6). Previous genetic studies identified CpgA-C and CCNA_01210 as key enzymes for CPG2 synthesis (6, 12). Deletion of *ccna_01210* resulted in the specific loss of CPG2 while retaining CPG (Fig. 1A) (12), suggesting that CCNA_01210 was involved in the conversion of CPG to CPG2. Complementation of the *ccna_01210* deletion restored CPG2 production (Fig. 1A) (12); we therefore refer to CCNA_01210 as CpgD for the remainder of this study. CpgD is annotated as a nucleotidyltransferase family protein. A structural homology search using the Alphafold predicted structure of CpgD (16) identified *Thermotoga maritima* inositol-1-phosphate cytidylyltransferase (IMPCT; PDB 4JD0) (17) as the top hit (Fig. 1B). Sequence alignment showed 24% identity and 40% similarity between IMPCT and CpgD, and three critical active site residues (R16, K26, and D112) were conserved in CpgD (Fig. 1C) (17). IMPCT uses CTP and inositol-1-phospate to produce CDP-inositol, an intermediate in di-myo-inositol-1,1’-phosphate (DIP) synthesis. Other members of this cytidylyltransferase family use phosphocholine (18) and phosphoglutamine (19) as substrates; we therefore hypothesized that CpgD may use phosphoglycerate to form CDP-glycerate.

**Figure 1:**
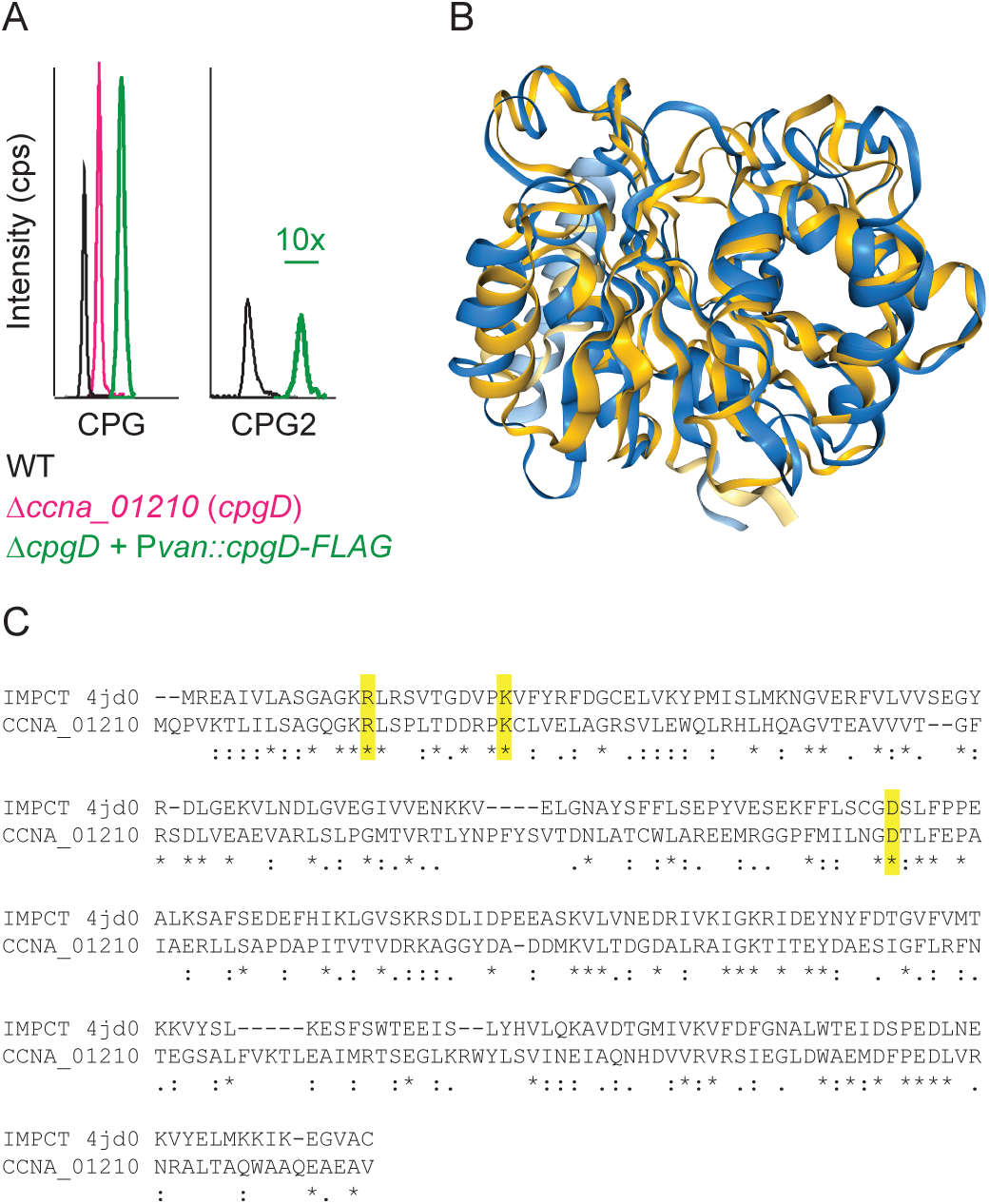
Identification of CpgD as a putative cytidylyltransferase. (A) Extracted-ion chromatograms demonstrate that the deletion of *ccna_01210* (*cpgD*) results in the complete loss of CPG2. CPG2 production is restored upon complementation of *cpgD*. Peaks are offset horizontally to enhance visibility. The CPG2 peak in the complementation strain is magnified 10x. (B) Foldseek (30) was used to generate an alignment of the Alphafold-predicted structure of CpgD (blue) with IMPCT (gold; PDB Accession 4JD0) (17). The proteins had an RMSD of 2.73 Å. (C) The sequence alignment of CpgD and IMPCT shows conservation of three active site residues highlighted in yellow.

### Purification and initial characterization of CpgD

We overexpressed and purified an N-terminal 6xHis-tagged CpgD from *E. coli* for biochemical analysis (Fig. 2A). To identify potential nucleotide substrates, we performed thermal shift assays in the presence of various nucleoside triphosphates; an increase in the melting temperature of the protein was only observed upon addition of CTP suggesting that CTP is the correct substrate (Fig. 2B). We incubated CpgD with CTP and 3-phosphoglycerate in the presence of Mg^2+^ and separated the nucleotide species by high-performance anion-exchange chromatography (HPAEC). When comparing the reaction chromatogram to those of CTP and CDP standards, we observed a new peak with a retention time of 2.7 minutes (Fig. 2C). This peak was absent in a control reaction containing no enzyme, suggesting that this was the product of CpgD. Mass spectrometry analysis of this peak was consistent with CDP-glycerate (whose [M-H]^-^ molecular ion is observed at *m/z* 490.027), and tandem MS/MS analysis confirmed the expected ion fragmentation (Fig. 2D).

**Figure 2:**
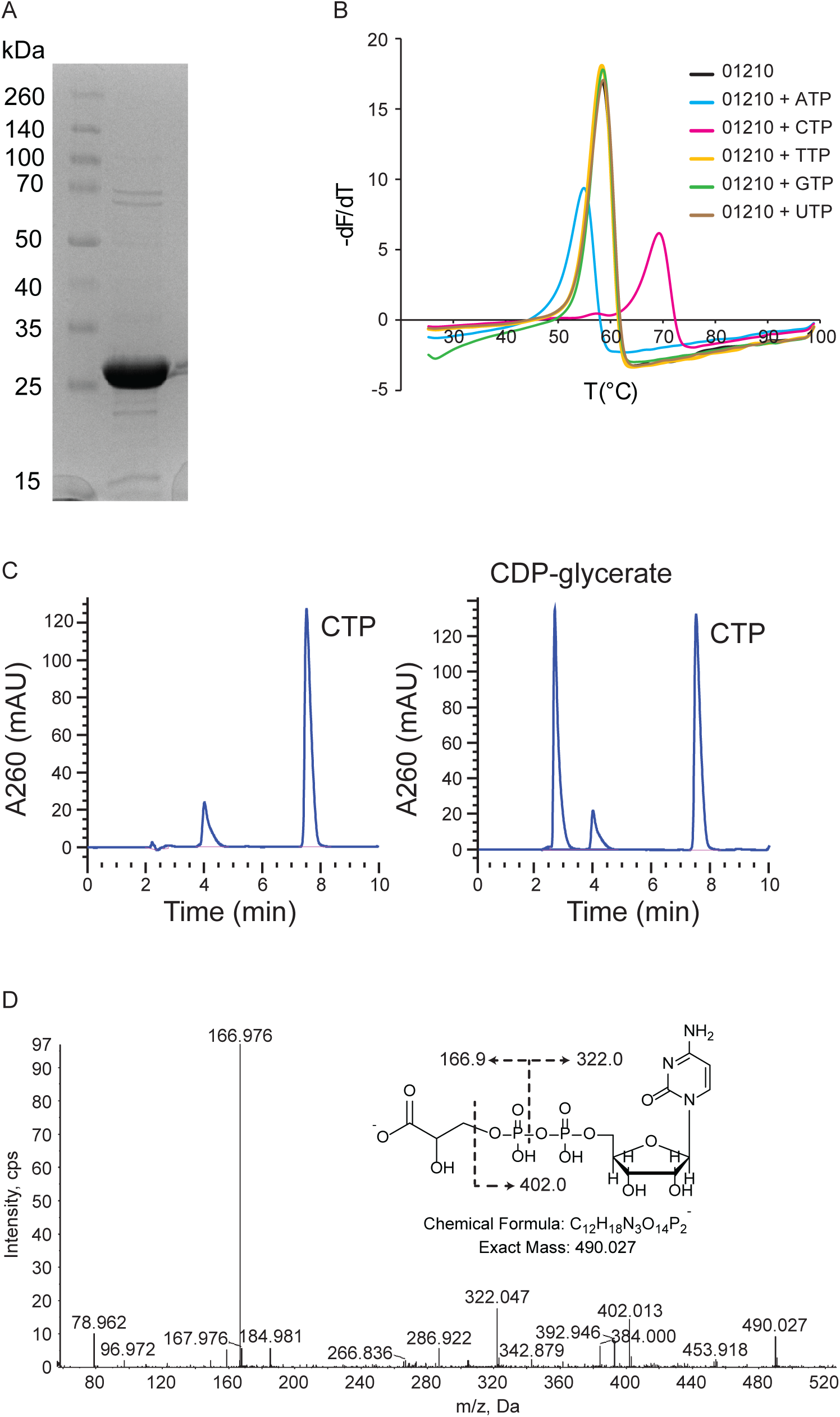
Initial characterization of CpgD activity. (A) His-tagged CpgD was purified from *E. coli*, resolved by SDS-PAGE, and visualized by Coomassie blue staining. (B) Thermal shift assays with the indicated nucleotides show an increase in Tm upon the addition of CTP. (C) HPAEC analysis of a CTP standard (left) or a reaction containing CpgD with CTP and 3-phosphoglycerate (right) shows the appearance of a peak in the reaction mixture with a retention time of 2.7 min. (D) The reaction mixture was analyzed by LC/ESI-MS/MS in the negative ion mode to determine the identity of the reaction product. The MS/MS fragmentation products are consistent with CDP-glycerate.

### Effect of pH and divalent cations on CpgD activity

To characterize the optimum conditions for CpgD activity, we quantified CDP-glycerate production over a pH range of 4.5-10; highest activity was seen between pH 7.5-8 (Fig. 3A). We next tested CpgD activity in the presence of various divalent cations (Fig. 3B). Highest activity was seen in the presence of magnesium with significant activity also observed in the presence of copper or cobalt. Lower activity was measured with zinc or manganese, and we did not observe any activity with calcium or in the absence of cations.

**Figure 3:**
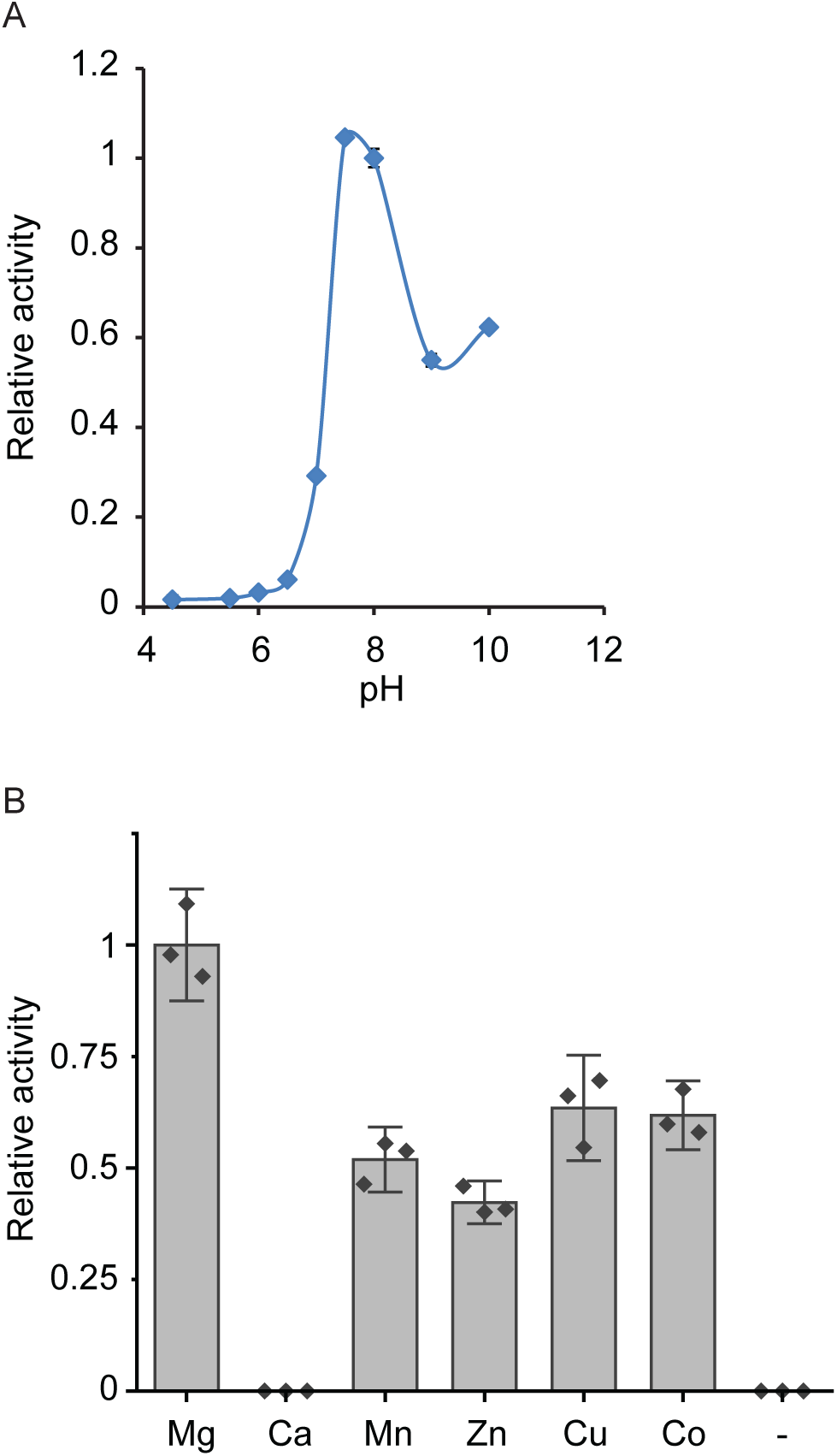
Characterization of CpgD pH and divalent cation requirements. (A) CpgD activity was determined over a range of pH’s using the following buffers: acetate (pH 4.5-5.5), HEPES (pH 6.5 – 8), Tris-HCl (pH 9), and borate (pH 10). Activity was quantified by HPAEC (n=3, error bars are the SD). (B) The activity of CpgD was determined in the presence of 50 mM of the indicated divalent cations. Activities were normalized to magnesium (n=3; error bars are the SD).

### Determination of CpgD kinetic parameters

Under the optimal conditions of pH 8 in the presence of Mg^2+^, we measured CDP-glycerate production over a period of 4 hours to determine the linear range of activity (Fig. 4A). Unless otherwise specified, all kinetic studies described below were performed for 3.5 hours in the presence of Mg^2+^ at pH 8. CpgD-catalyzed production of CDP-glycerate exhibited typical Michaelis–Menten kinetics with respect to 3-phosphoglycerate (Km,app = 10.9 ± 0.7 mM; Vmax,app = 0.72 ± 0.02 µmol/min/mg enzyme) and CTP (Km,app = 4.8 ± 0.9 mM; Vmax,app = 0.44 ± 0.03 µmol/min/mg enzyme) (Fig. 4C-D). These Km values are consistent with the measured intracellular concentrations of CTP (2.7 mM) and 3-phosphoglycerate (1.5 mM) in *E. coli* (20).

**Figure 4:**
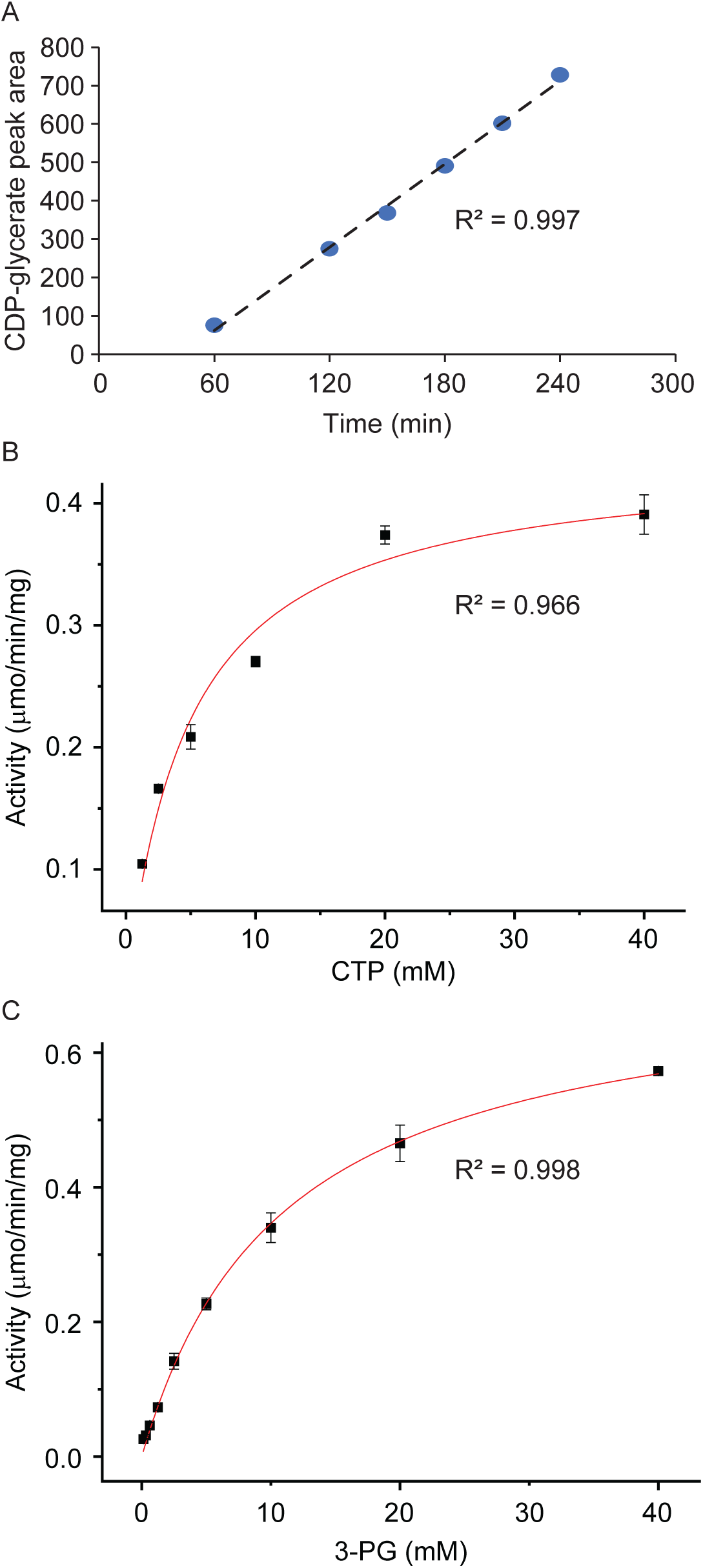
CpgD enzyme kinetics. The kinetic parameters of CpgD were measured by HPAEC. (A) CpgD activity was measured as a function of time to ensure reactions were assayed within the linear range of velocity. (B-C) Michaelis-Menten kinetic parameters were determined for CpgD (n=3, error bars are SD). (B) To determine the K_m,app_ for CTP, 3-phosphoglycerate concentration was held constant (10 mM) while CTP concentration varied. (C) The K_m,app_ for 3-phosphoglycerate was determined by holding the CTP constant at 10 mM while varying the 3-phosphoglycerate concentration. K_m,app_ values were 4.8 ± 0.9 mM and 10.9 ± 0.7 mM for CTP and 3-phosphoglycerate, respectively. V_max,app_ values were 0.44 ± 0.03 µmol/min/mg enzyme and 0.72 ± 0.02 µmol/min/mg enzyme for CTP and 3-phosphoglycerate, respectively.

## Discussion

Bacteria produce sphingolipids with diverse headgroups including sugars (10), phosphoglycerol (21), phosphoglycerate (6), and phosphoethanolamine (7). The CPG and CPG2 lipids help enable *C. crescentus* survival in the absence of LPS (6). Of the genes identified to play a role in CPG2 synthesis (6, 12), the only one with an ascribed function is the ceramide kinase, CpgB (13). Here, we show that CCNA_01210 (CpgD) is a cytidylyltransferase which produces CDP-glycerate, a required metabolite for CPG2 synthesis.

The wall teichoic acid (WTA) synthesis pathway in Gram-positive bacteria uses a similar metabolite, CDP-glycerol. These organisms use the COG0615-domain cytidylyltransferases (22) TagD or TarD to produce CDP-glycerol from CTP and phosphoglycerol (23, 24). Interestingly, despite having similar substrates and products, TagD/TarD and CpgD have no homology. Instead, CpgD has sequence and predicted structural homology to cytidylyltransferases containing the COG1213 conserved domain (22) (Fig. 1B-C). COG1213-domain cytidylyltransferases have been reported to use phosphosugars, phosphocholine, and phoshoglutamine as substrates (16, 18, 19). Although we are not aware of any COG1213 enzymes that use phosphoglycerate as a substrate *in vivo*, the phosphoglutamine cytidylyltransferase from *Campylobacter jejuni* (Cj1416) exhibits high substrate promiscuity and can use phosphoglycerate *in vitro* when incubated with manganese rather than magnesium (19). Further structural characterization of CpgD will provide insight into the mechanism of substrate specificity among these enzymes.

Our characterization of CpgD solves one more piece of the puzzle to the enzymatic pathway responsible for CPG2 synthesis (Fig. 5). Future work will be required to determine the activities of CpgA/C as well as any other, yet unidentified, enzymes.

**Figure 5:**
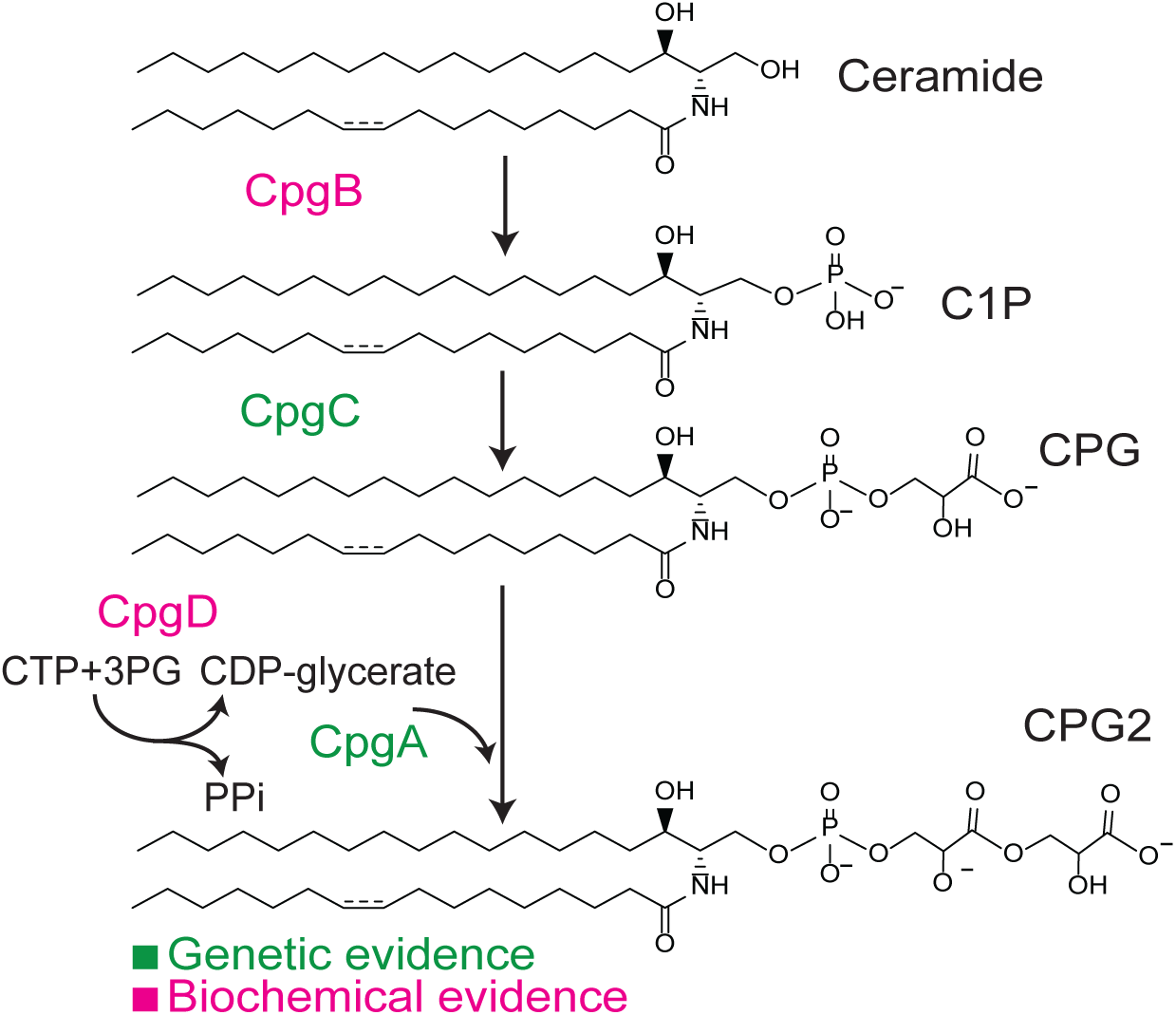
Proposed synthetic mechanism for CPG2. Based on previous genetic (6, 12) and biochemical (13) studies, we propose the following pathway for CPG2 synthesis. CpgB is a ceramide kinase that uses ATP and ceramide to produce ceramide 1-phosphate (C1P). CgpC, through a yet unidentified mechanism, converts C1P to CPG. CpgD uses CTP and 3-phosphoglycerate to produce CDP-glycerate, which is then a substrate for CpgA to convert CPG to CPG2. Steps with biochemical evidence are highlighted in magenta and those with genetic evidence are highlighted in green.

## Experimental Procedures

### C. crescentus growth conditions

*C. crescentus* wild-type strain NA1000, and its derivatives were grown at 30 °C in peptone-yeast-extract (PYE) medium (25) for routine culturing. When necessary, antibiotics were added at the following concentrations: kanamycin 5 µg/ml in broth and 25 µg/ml in agar (abbreviated 5:25); spectinomycin 25:100. Gene expression was induced in *C. crescentus* with 0.5 mM vanillate.

### Deletion and complementation of ccna_01210 (cpgD)

The primers used for cloning are listed in Table 1. Δ*cpgD* was cloned by PCR amplifying the upstream (EK1698/1699) and downstream (EK1700/1701) homology fragments from NA1000 genomic DNA. The fragments were stitched together by overlap PCR. The final purified PCR product was ligated into the EcoRI/HindIII site of pNPTS138 (M.R.K. Alley, unpublished). The assembled plasmid was electroporated into NA1000 followed by selection on PYE-kanamycin plates. An individual colony was grown overnight in PYE and streaked onto PYE-3% sucrose plates for *sacB* selection. Colonies were screened for the *cpgD* deletion by PCR (EK S355/S356; wild-type 1.8 kb; deletion 1.4 kb). Flag-tagged *cpgD* was amplified from NA1000 genomic DNA (EK1740/1741). The PCR product was ligated into the NdeI/NheI site of pVCFPC-1 (26). The resulting plasmid was electroporated into the Δ*cpgD* strain.

**Table 1:**
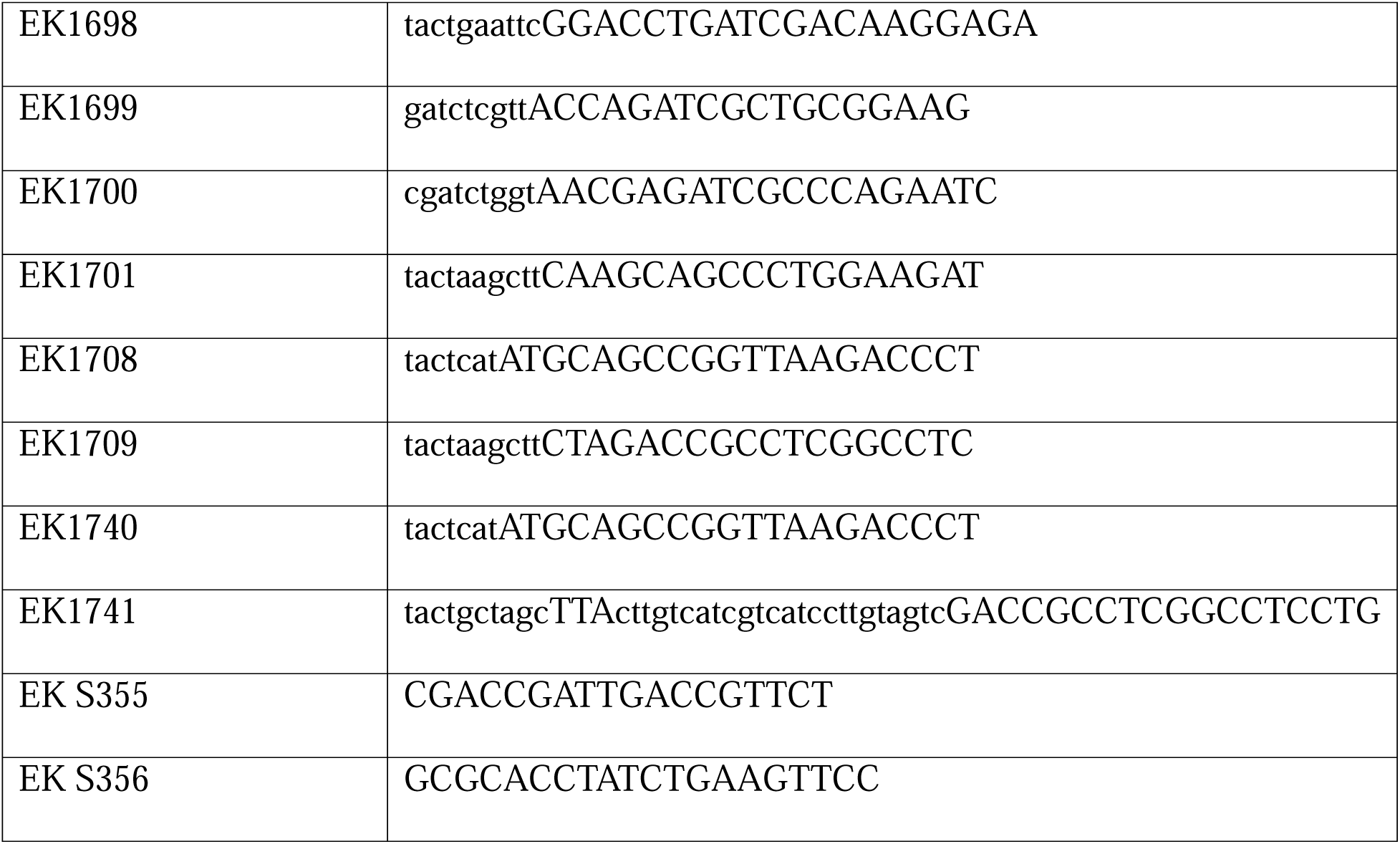
Primers used in this study.

### Lipidomic profiling and confirmation of ceramide-phosphate production by LC/MS/MS

Lipids were extracted from *C. crescentus* cells using the method of Bligh and Dyer with minor modifications (27). The lipid extracts were analyzed by normal phase LC/MS/MS in the negative ion mode as previously described (28, 29).

### Purification of His-tagged CpgD

*cpgD* was amplified (EK1708/1709) and ligated into the NdeI/HindIII site of pET-28a (EMD Millipore) to yield an N-terminal fusion protein. The resulting plasmid was transformed into *E. coli* strain ER2566 (New England Biolabs) for expression and purification. A 1 L culture of *E. coli* ER2566 cells carrying the pET28a-*cpgD* plasmid was grown in LB broth with 30 μg/ml kanamycin at 37 °C with shaking until reaching an A_600_ of 0.3. IPTG was added to a final concentration of 0.25 mM, followed by induction at 30 °C for 3 h. Cells were harvested by centrifugation at 10,000 x g. The pellet was resuspended in 25 ml of lysis buffer (50 mM NaH_2_PO_4,_ 300 mM NaCl, 10 mM imidazole) before lysing via 2 to 3 passages through a French press (11,000 psi). The lysate was centrifuged at 8000 x g for 10 min to remove unbroken cells and the supernatant was passed through a 0.22 mm syringe filter. The His-tagged CpgD was purified using an ÄKTA start FPLC system and a 1 ml HisTrap HP column (Cytiva). Once the sample was loaded it was washed with lysis buffer to remove the unbound material. Elution was carried out via a linear gradient of 10-1000 mM imidazole. 1 ml fractions were collected, and the protein elution was monitored by A_280_. Fractions which contained the purified protein were identified by SDS-PAGE and Coomassie blue staining. Fractions containing the protein were pooled and dialyzed into a buffer containing 10 mM Tris pH 8.0, 0.1 M NaCl, 2 mM EDTA and 1 mM DTT over 36 h at 4 °C using a Slide-A-Lyzer Dialysis Cassette with a 20,000 MW cutoff (Thermo Scientific). The protein concentration was determined using the BCA Protein Assay Kit (Pierce).

### Thermal shift protein stability assay of CpgD

A 20 ml reaction was set up containing 1X GloMelt dye (Biotium), 1.2 mg/ml CpgD, 50 mM Tris pH 8.0, 50 mM MgCl_2_, 40 nM ROX, and 10 mM nucleotide. Samples without nucleotide were used as a negative control. Reactions were carried out in triplicate and the melt curve profile was assayed on an ABI QuantStudio 6 Flex (Thermo Fisher Scientific). Initial hold was at 25 °C for 30 s and the melt curve was measured between 25-99 °C with a ramp rate of 0.05 °C/s. Data were plotted using first derivative (slope) of the fluorescence curve against temperature to calculate the Tm for each sample.

### CpgD activity assay

Enzyme activity assays for CpgD were carried out in a total volume of 20 µl containing 50 mM Tris pH 8.0, 50 mM MgCl_2_, 10 mM CTP, and 10 mM 3-phosphoglycerate. The reaction was initiated with the addition of 0.24 mg/ml CpgD and proceeded at 35 °C for 3.5 h. Reactions were stopped by heating at 90 °C for 5 minutes and stored at -20 °C until analysis.

### Detection of products using High Performance Anion-Exchange Chromatography (HPAEC)

The chromatography apparatus included an Agilent Technologies 1200 series HPLC equipped with a quaternary pump (G5611A), Infinity Bio-Inert HPLC Autosampler (G5667A), MWD Detector (G1365C), and OpenLAB Control Panel Software (version A.01.05). Samples were resolved on a Dionex CarboPac PA200 column (3 mm×250 mm) with the corresponding guard column (3 mm×50 mm) (Thermo Fisher). The column was equilibrated with 60% Buffer A (1 mM NaOH) and 40% Buffer B (1 M sodium acetate in 1 mM NaOH), and the column temperature was maintained at 30 °C. Samples were diluted 1:100 in Milli-Q water and 25 µl was injected for HPAEC. The HPAEC run consisted of a 10 min linear gradient from 60:40 to 20:80 Buffer A: Buffer B, 2 min hold at 100% Buffer B, and 5 min of 60:40 of Buffer A: Buffer B, to re-equilibrate the column. The entire run was carried out at a flow rate of 0.4 ml/min. CTP and CDP-glycerate were monitored by their absorbance at 260 nm. Product formation was measured by calculating peak area using OpenLAB (version A.01.05). A standard curve of CTP was used to quantify product formation.

### Identification of CpgD product

A reaction containing 50 mM Tris pH 8.0, 50 mM MgCl_2_, 10 mM CTP, 10 mM 3-phosphoglycerate, and 0.24 mg/ml CpgD was incubated at 35 °C for 3.5 h. The reactions were stopped by the addition of methanol (4x reaction volume), and samples were centrifuged at 10,000 rpm at room temperature for 2 min. Subsequently, the supernatant was collected and used for LC/MS/MS analysis.

#### LC/MS/MS

Reverse-phase liquid chromatography-electrospray ionization/tandem mass spectrometry (LC-ESI/MS/MS) was performed using a Shimadzu LC system (comprising a solvent degasser, two LC-10A pumps and a SCL-10A system controller) coupled to a high-resolution TripleTOF5600 mass spectrometer (AB Sciex, Framingham, MA). LC was operated at a flow rate of 200 μl/min with a linear gradient as follows: 100% of mobile phase A was held isocratically for 2 min and then linearly increased to 100% mobile phase B over 5 min and held at 100% B for 2 min. Mobile phase A was a mixture of water/acetonitrile (98/2, v/v) containing 0.1% acetic acid. Mobile phase B was a mixture of water/acetonitrile (10/90, v/v) containing 0.1% acetic acid. A Zorbax SB-C8 reversed-phase column (5 μm, 2.1 x 50 mm) was obtained from Agilent (Palo Alto, CA). The LC eluent was introduced into the ESI source of mass spectrometer. Instrument settings for negative ion ESI/MS and MS/MS analysis of lipid species were as follows: Ion spray voltage (IS) = -4500 V; Curtain gas (CUR) = 20 psi; Ion source gas 1 (GS1) = 20 psi; De-clustering potential (DP) = -55 V; Focusing potential (FP) = -150 V. Data acquisition and analysis were performed using the Analyst TF1.5 software (AB Sciex, Framingham, MA).

### Determining the optimum pH conditions for CpgD activity

Enzyme activity assays were performed as above with the pH adjusted to 4.5, 5.5, 6, 6.5, 7, 7.5, 8, 9, or 10. The pH was adjusted using the following buffers: acetate (pH 4.5 – 5.5), HEPES (pH 6.5 – 8), Tris-HCl (pH 9), and borate (pH 10). Enzyme activities were normalized to the activity at pH 8.0.

### Characterization of the divalent cation requirement for CpgD activity

Activity assays were performed as above using 50 mM of the following: magnesium chloride, manganese sulfate, calcium chloride, zinc sulfate, copper sulfate, or cobalt nitrate. A control reaction was set up with no metal ions. Activities were normalized to that observed with MgCl_2_.

### CpgD kinetic analyses

For kinetic analyses, reactions were performed as mentioned above while varying the substrate concentrations. To determine the K_m,app_ for 3-phosphoglycerate, CTP concentration was held constant (10 mM) while 3-phosphoglycerate concentration ranged from 0.125 to 40 mM. The K_m,app_ for CTP was determined by holding the 3-phosphoglycerate constant at 10 mM while varying the CTP concentration from 1.25 to 40 mM. The enzyme activity was fit to the Michaelis–Menten equation using OriginPro (OriginLab).

## Data availability

All of the data for this work is contained within the manuscript.

## Author CrediT statement

**Tanisha Dhakephalkar:** Conceptualization, Methodology, Investigation, Writing-Original Draft, Visualization. **Ziqiang Guan:** Conceptualization, Investigation, Writing-Review & Editing. **Eric Klein:** Conceptualization, Methodology, Writing-Original Draft, Visualization, Supervision.

